# The anterior cruciate ligament in murine post-traumatic osteoarthritis: markers and mechanics

**DOI:** 10.1101/2021.11.08.467455

**Authors:** Lorenzo Ramos-Mucci, Ahmed Elsheikh, Craig Keenan, Ashkan Eliasy, Kris D’Aout, George Bou-Gharios, Eithne Comerford, Blandine Poulet

**Author notes:** **Correspondence:** Dr Blandine Poulet, Institute of Life Course and Medical Sciences, University of Liverpool, Apex building, West Derby Street, Liverpool, L7 8TX, UK.

## Abstract

Osteoarthritis (OA) is a whole joint disease that affects all knee joint tissues. Ligaments, a matrix-rich connective tissue, play an important mechanical function that stabilises the knee joint and yet their role in OA is not well studied. Recent studies have shown that ligament extracellular matrix (ECM) structure is compromised in the early stages of OA, but it remains unclear how this affects ligament function and biomechanics. In this study, the aim was to investigate the structural, cellular and viscoelastic changes in the anterior cruciate ligament (ACL) in a murine non-invasive post-traumatic OA (PTOA) model. Non-invasive mechanical loading of the knee joint of C57BL/6J mice (10-week-old) was used as a PTOA model. Knee joints were analysed for joint space mineralisation and the ACLs were assessed with histology and mechanical testing. PTOA knee joints had a 33-46% increase in joint space mineralisation and PTOA knee joint ACLs exhibited ECM modifications, including collagen birefringence and COL2 and proteoglycan deposition. ECM changes were associated with cells expressing chondrogenic markers (SOX9 and RUNX2) expanding from the tibial enthesis to the ACL midbody. Viscoelastic and mechanical changes in the ACLs from PTOA knee joints included a 20-21% decrease in tangent modulus at 2MPa of stress, and a decrease strain rate sensitivity at higher strain rates and a significant increase in relaxation during stress-relaxation, but no changes to hysteresis and ultimate load to failure. These results demonstrate that ACL pathology and viscoelastic function is compromised in murine PTOA knee joints and provides further evidence of the important role of ligaments in the knee joint organ in health and disease.

## 2 Introduction

Osteoarthritis (OA) is a degenerative joint disease that affects the whole knee joint (Loeser *et al.*, 2012), including loss of articular cartilage, osteophyte formation, subchondral bone remodelling, synovial hyperplasia and ligament degeneration. Ligaments are the main stabilizers of the knee joint (Frank, 2004), however their role in OA progression is largely unexplored. Previously, we showed structural and pathological changes in the cruciate and collateral knee ligaments of murine OA mouse models (Ramos-Mucci *et al.*, 2020), similar to the pathology described in human OA anterior cruciate ligaments (ACLs) (Hasegawa *et al.*, 2013; Hasegawa *et al.*, 2012). These structural and pathological changes could alter the ligament mechanical function and ultimately induce further OA.

Tissue structure and mechanical function are closely related, including in ligaments (Frank, 2004). Ligaments are composed of a collagen hierarchical structure made of primarily collagen type I as well as noncollagenous extracellular matrix (ECM) proteins such as proteoglycans, elastin and glycolipids (Frank, 2004). Ligaments are viscoelastic and demonstrate viscoelastic properties such as: 1) strain rate sensitivity, where mechanical behaviour is stiffer at higher strain rates, 2) stress-relaxation, where stress reduces under constant deformation, 3) creep, where deformation occurs under a constant load and 4) hysteresis, where there are differences in the loading and unloading behaviour (Woo, 1982). Viscoelastic behavior is mainly attributed to collagen interaction with ECM structural proteins such as proteoglycans and elastin (Elliott *et al.*, 2003; Franchi *et al.*, 2007; Robinson *et al.*, 2004). Specifically, ECM changes that affected viscoelastic properties of ligaments or tendons include collagen (Connizzo *et al.*, 2015; Sun *et al.*, 2015), proteoglycans (Connizzo *et al.*, 2013) and elastin content (Henninger *et al.*, 2015).

The consequences of the OA disease process on the mechanical behaviour of ligaments are not fully understood. To date, there are few studies that characterise the mechanical properties of the ACL in OA knee joints. In human knee OA patients, one study found a decrease in elastic stiffness and an increase in viscoelastic stress-relaxation in ACLs (Hagena *et al.*, 1989). In the Dunkin Hartley guinea pig spontaneous OA model, there were reduction in the viscoelastic toe-region laxity of the ACLs compared to healthy controls (Quasnichka *et al.*, 2005). In STR/ort spontaneous OA mice, ACLs had a lower ultimate load compared to healthy control knee joints (Anderson-MacKenzie *et al.*, 1999). These studies confirm a decrease in stiffness and ultimate load in the ACL of OA knee joints but did not measure other physiologically relevant viscoelastic properties of the ACL (Franchi *et al.*, 2007).

Murine models offer several advantages to study OA progression. The post-traumatic OA (PTOA) model developed by Poulet et al. offers the ability to test the ACL mechanical properties post-trauma without invasive ligament transection (Poulet *et al.*, 2011). The PTOA mouse knee joint is known to develop OA in the lateral knee compartment with moderate severity following two weeks of non-invasive loading of the knee joint (Poulet *et al.*, 2011). The pathological and mechanical properties of the ACL have not previously been studied in the PTOA knee joint.

There is a lack of knowledge of the ligament cellular microenvironment and its surrounding structural and mechanical niche. Understanding these components can help us understand biomechanical pathways in the ligaments driving ligament degeneration and potentially knee joint OA progression. We hypothesise that changes in the ACL ECM composition and structure during OA development will result in reduced stiffness and modifications of their viscoelastic properties. The aim of this study was to characterise the ACL mechanical, matrix and cellular environment during murine PTOA pathology, to better understand PTOA disease processes for future therapeutical interventions.

## 3 Materials and Methods

### 3.1 Post-traumatic osteoarthritis (PTOA) murine model

Non-invasive joint loading was performed on the right knees of ten-week-old C57BL/6 mice (Charles River, UK) to induce OA and articular cartilage lesions as has been previously shown (Poulet *et al.*, 2011). Briefly, axial 9N compressive loads were applied for forty cycles in each loading episode, and six loading episodes over a period of two weeks. Mice were kept under general gas anaesthesia (isoflurane) for the entire procedure, which lasts about 10 minutes.

For joint space mineralisation quantification, knee joints were analysed at 4 (n=7) and 14 (n=4) weeks after the last loading episode, to determine progressive changes following OA induction. C57/BL6 male mice at 14 and 24 weeks of age were used as age-matched controls (n=4 per group).

For ACL histology and mechanical testing, knee joints were analysed six weeks after the last loading episode. From the PTOA mice (n=12 total); n=8 of the right loaded legs were used for ACL mechanical testing, and n=4 right loaded legs for μCT (micro-computed tomography) ACL measurements. From the non-loaded mice, right legs were used for ACL mechanical testing (n=8), and left legs for μCT ACL measurements (n=4). Left knee joints were used to measure ACL measurements, as it has been previously shown that contralateral ACLs are an appropriate surrogate to for anatomical dimensions (Jamison *et al.*, 2010).

All mice were kept in the same conditions in polypropylene cages, subjected to 12-hour light/dark cycles, at 21±2°C and fed standard diet ad libitum. For tissue collection, animals were euthanised by cervical dislocation. All procedures were performed in accordance with the UK Home Office guidelines and regulations under the Animals (Scientific Procedures) Act 1986 and local ethics committee (project license: P267B91C3).

### 3.2 Measuring knee joint space mineralisation in the murine PTOA knee joint

Knee joint mineralisation was quantified as previously described (Ramos-Mucci *et al.*, 2020). PTOA knee joint were analysed at 4 and 14-weeks post-trauma. Briefly, cadaveric knee joints were fixed (neutral buffered formalin) and dehydrated (in 70% ethanol) and then were scanned using a Skyscan 1172 μCT scanner (Skyscan, Belgium) with a 5μm isotropic voxel size (50kV, 200μA respectively, 0.5mm Aluminium filter; 0.6° rotation angle, no frame averaging). Regions of interests of the knee joint comprising of menisci and ligaments were hand-drawn and quantified using CTAn (Skyscan, Belgium) (Ramos-Mucci *et al.*, 2020). Statistical analysis of mineralised volume of the murine knee joint was performed using a one-way ANOVA and post-hoc analysis to compare control and PTOA knee joints at 4 and 14-weeks post-trauma (significance was set at p < 0.05). Three-dimensional models of the mineralised tissue were created using CTVox from the region of interest selected for mineralised tissue volume analysis (Skyscan, Belgium).

### 3.3 Histological and immunohistochemical assessment of the murine knee joint

For histology, fixed (neutral buffered formalin) joints were decalcified (ImmunocalTM, Quarttet, Berlin, Germany), dehydrated and processed for wax embedding. Serial coronal 6μm thick sections were cut across the entire joint. Slides were stained with toluidine blue (TB) to assess pathophysiological changes in the knee joint (Glasson *et al.*, 2010) and picrosirius red to assess collagen birefringence (Junqueira *et al.*, 1979). TB staining consisted of 0.1% TB in 0.1M solution of acetate buffer, pH 5.6 for 15 minutes and counterstained with 0.2% fast green for 5 seconds and then rinsed with acetone twice for 10 seconds. For picrosirius red, sections were stained in Weigert’s haematoxylin for 8 minutes then stained with picrosirius red solution (0.1% w/v sirius red in saturated aqueous picric acid) for one hour and then washed with 0.5% v/v glacial acetic acid in distilled water for 3 minutes. Slides were then dehydrated, mounted and imaged with an Olympus microscope with an adjustable polariser (Olympus BX60).

Immunohistochemistry was performed to localise expression of collagen type II (COL2) (Thermo, Mouse MC), SOX9 (Millipore, Rabbit PC), RUNX2 (Abcam, Rabbit PC) and asporin (ASPN) (Abcam, Rabbit PC), as described previously (Ramos-Mucci *et al.*, 2020). For COL2, antigen retrieval was applied with pepsin (3mg/mL in 0.02M HCl) for 45 minutes at 37ᵒC. Slides were then blocked for endogenous peroxidase with 0.3% hydrogen peroxide (Sigma, 15 minutes), for endogenous Avidin/Biotin binding with an Aviding/Biotin Blocking Kit (Vector Labs, SP2001; 15 minutes each), and for non-specific binding sites (COL2: blocking solution from MoM Kit, Vector Labs, BMK-2202; SOX9, RUNX2, ASPN: 10% v/v goat serum). Primary antibodies were incubated overnight at 4°C and included COL2 (1/100), SOX9 (1/1000), RUNX2 (1/1000), ASPN (1/400). Negative controls included a mouse IgG (2μg/mL, Sigma) or rabbit IgG (1μg/mL, Vector Labs). Biotinylated secondary antibody (Vector Labs; 1/200 for 1h) and then Vectastain solution (Vector Labs) were applied. Stains were developed with DAB (Vector Labs), dehydrated, mounted in DPX and imaged with a Zeiss microscope (Zeiss).

### 3.4 Murine anterior cruciate ligament (ACL) imaging and measurements

Mouse knee joints were analysed with μCT to quantify the murine ACL length and cross-sectional area (CSA). PTOA knee joint ACL samples from the right loaded knee joint were used (n=4), and in the healthy control group ACL samples from the left knee joint were used (n=4). Low sample number of ligaments is a potential limitation of this method. Immediately after death, skin was removed from the hind limb and was frozen and kept at −20℃ until analysis. The knee joints were defrosted and dissected under a microscope until only the femur-ACL-tibia complex remained. Femur-ACL-tibia samples were submerged in Lipiodol contrast agent (Guerbet, FR) overnight at 4℃. Sample was fitted into a small straw holder filled with Lipiodol and a putty adhesive (Blu Tack) was used to keep the femur in place at a 90° angle with the ACL and tibia suspended vertically. The femur-ACL-tibia samples were then scanned with a 5μm isotropic voxel size (50kV, 200μA respectively, 0.5mm Aluminium filter; 0.6° rotation angle, no frame averaging) using a Skyscan 1172 μCT scanner (Bruker, BE). This is similar to previous methods using contrast agents to image the mouse knee joint (Das Neves Borges *et al.*, 2014).

A sagittal area of interest, showing the femoral condyles-ACL-and tibia condyles, was used to measure ACL length using the CTan software (Bruker) (Fig. 1A, yellow arrow). A second area of interest of the entire ACL volume was selected in the Dataviewer software (Bruker) and saved in the axial plane for analysis of the ACL CSA. The ACL CSA was measured using the CTan software (Bruker) at the ACL midsection (Fig. 1A, yellow arrow). An average ACL length and CSA was calculated for each mouse group and these were used to calculate stress-strain.

**Figure 1:**
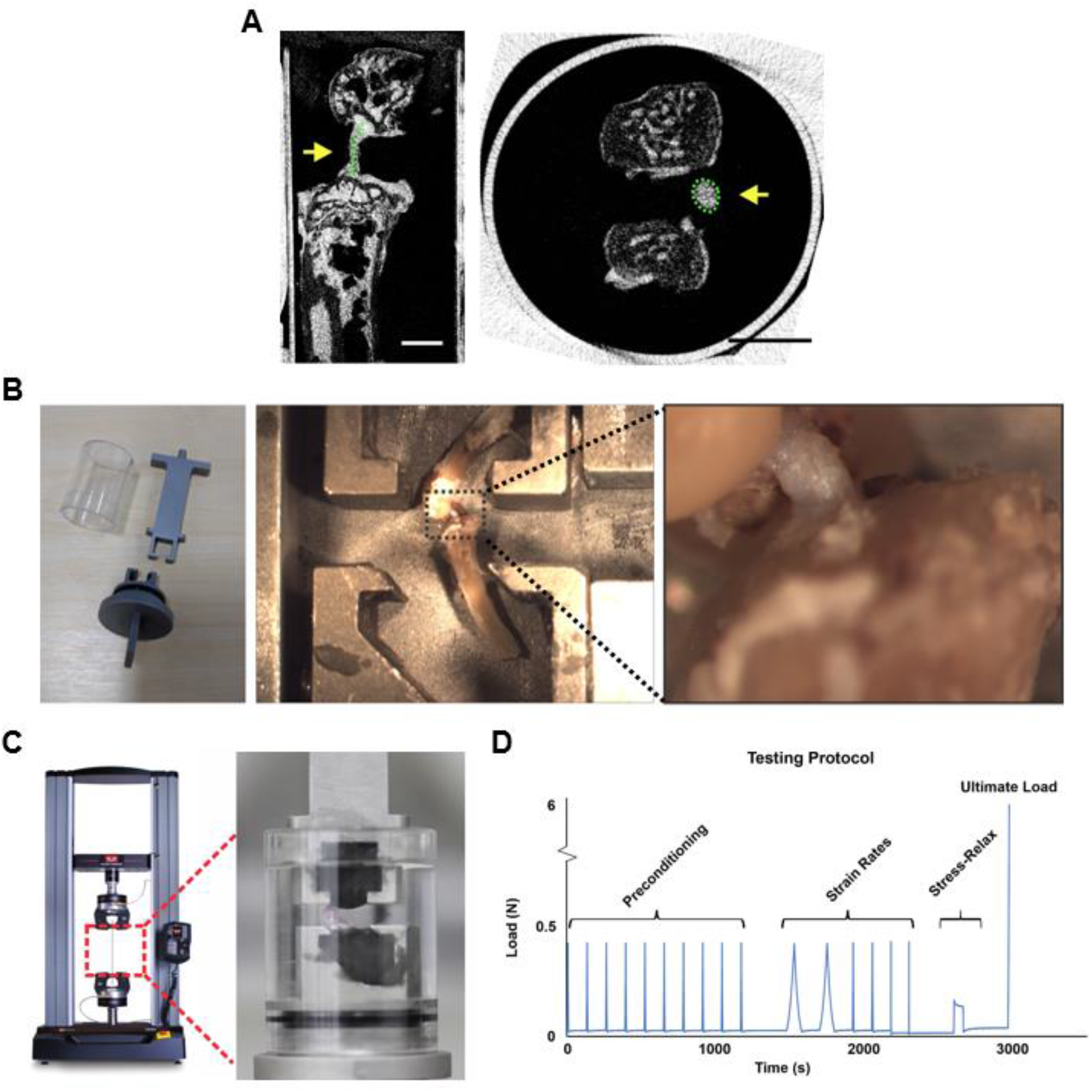
Murine anterior cruciate ligament (ACL) imaging and mechanical testing setup. A) Murine ACL was visualised with μCT imaging with a contrast agent and used to measure length and cross-sectional area (CSA) (yellow arrows). B) Mechanical testing setup included custom-made clamps, which allowed for poly(methyl methacrylate) fixation of the femur and tibia at a 90° angle and for vertical alignment of the ACL. C) Fixed samples were loaded into a dual column uniaxial machine (Instron) for mechanical testing of the ACL. D) Mechanical testing protocol used for viscoelastic and material behaviour of the ACL which included preconditioning cycles, strain rate testing at 0.1%/s, 1%/s and 10%/s strain rates, stress-relaxation loaded to 5% strain and lastly ultimate load to failure was tested at 1%/s until ACL rupture.

### 3.5 Mechanical testing of the murine anterior cruciate ligament (ACL)

Murine ACL mechanical properties were analysed with a customised clamp and tensile testing system (Fig. 1B), designed based on past literature (Anderson-MacKenzie *et al.*, 1999; Connizzo *et al.*, 2015; Sun *et al.*, 2015; Warden *et al.*, 2006). The clamp was manufactured in steel at the School of Engineering at the University of Liverpool.

Immediately following death, murine hind legs were stored at −20ᵒC. Storage at this temperature has been previously shown to not influence the viscoelastic and mechanical properties of ligaments (Moon et al., 2006; Woo et al., 1986). On the day of measurement, knee joints were defrosted and dissected until only the femur-ACL-tibia complex remained. Poly(methyl methacrylate) (PMMA) (Technovit 6091) was used to fix the femur and tibia, to approximate 90° of knee flexion (Fig. 1B). This extent of knee flexion was within the physiological range of mouse knee flexion (Sup. Table 1) and allowed for uniform tensile testing along the axis of the ligament (Warden *et al.*, 2006).

The femur-ACL-tibia was briefly aligned under the dissection microscope to ensure a vertical ACL orientation (Fig. 1B). The fixture with the sample was transferred to a dual column uniaxial material testing machine (Instron 3366, USA) with a 10N load cell. The encasing outer covering enclosed the sample and PBS was added to ensure a hydrated ligament sample (Fig. 1C).

The mechanical testing protocol applied a preload of 0.02N to remove laxity in the ligament, followed by a series of 10 preconditioning cycles at a strain rate of 1%/s to a maximum load of 0.4N, which ensured ligaments were in a steady state condition (Fung, 1993) (Fig. 1D). Following preconditioning, strain rate sensitivity, stress-relaxation behaviour and ultimate load at failure were tested (Fig. 1D). For strain rate testing, two load-unload cycles of three different strain rates were applied (0.1%/s, 1%/s, and 10%/s) (Fig. 1D). These strain rates have been used in previous studies on murine ACLs (Anderson-MacKenzie *et al.*, 1999; Sun *et al.*, 2015). Viscoelastic stress-relaxation was tested at 5% strain (applied strain at 1%/s) and elongation was maintained while monitoring load, similar to a previous study (Connizzo *et al.*, 2015). Elongation was maintained for 120 seconds where stress-relaxation behaviour started to stabilise (Fig. 1D). Lastly the ultimate load at failure was tested by increasing the load on the specimen while maintaining a strain rate of 1%/s until ligament rupture took place (Fig. 1D).

Analyses of the load-elongation data were performed on Microsoft Excel (version 2109). Stress, strain tangent modulus and hysteresis values were calculated as previously described (Elsheikh *et al.*, 2008; Readioff *et al.*, 2020). Average ACL length and CSA measurements were used for all these calculations (Section 3.4 and Sup. Table 2).

For stress-strain behaviour, the ACL length and CSA measurements for the PTOA and control ACL were used to calculate the stress and strain using Equations 1 and 2, respectively. The resulting stress-strain curve was then fitted to an exponential curve suitable for ligament stress-strain behaviour (Haut and Little, 1969) using the least squares method. From the exponential best-fit curve for each strain rate, the tangent modulus (gradient of the stress-strain curve) at each stress or strain was derived using Equation 3.

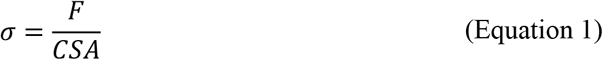

where σ is stress (MPa), F is the applied force (N) and CSA (mm^2^) is the estimated cross-sectional area.

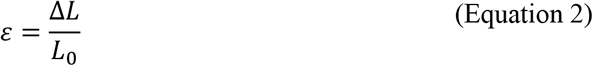

where ε is strain (mm/mm), ΔL is the corresponding change in elongation (mm) (ΔL = L_t_ – L_0_), L_0_ is the initial estimated ligament length (mm), and L_t_ is the deformed length of the ligament (mm).

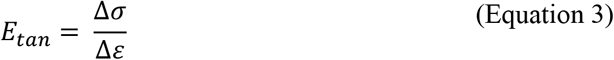

where E_tan_ is tangent modulus (MPa).

To determine the sensitivity of stress-strain behaviour to changes in strain rate, estimations of the stress-strain behaviour and tangent modulus were repeated for all the different strain rates (0.1%/s, 1%/s, 10%/s). Tangent modulus values were normalised to the 0.1%/s strain rate tangent modulus of the same sample using Equation 4.

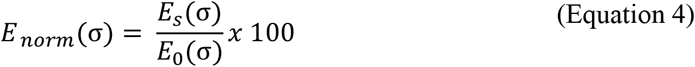

where E_norm_(σ) is the normalised tangent modulus (%) at a specific stress, E_tan_(σ) is the tangent modulus at 1%/s or 10%/s strain rates at the corresponding stress, and E_0_(σ) is the tangent modulus at 0.1%/s strain rate at the corresponding stress.

Stress-relaxation behaviour was characterised by monitoring the stress degradation over time while maintaining a strain of 5%. Stress was normalised by the peak stress at t=0 (when 5% strain was reached) (Connizzo *et al.*, 2015) using Equation 5. The stress-relaxation curve was fitted into a polynomial equation to determine the average normalised stress for all samples.

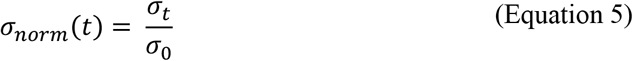

where σ_norm_(t) is the normalised stress (MPa/MPa) over log time, σ_o_ the peak stress at t=0 and σ_t_ is the stress behaviour over log time.

Hysteresis was calculated from the load-elongation data to measure the area between the load and unload cycles which correlates to the energy dissipated due to material viscosity. Hysteresis was measured using numerical integration (trapezoidal rule) as described previously (Elsheikh *et al.*, 2008; Readioff *et al.*, 2020).

### 3.6 Statistical analysis

For statistical analysis of the ACL viscoelastic properties (stress-strain, tangent modulus, strain rate sensitivity, normalised strain rate sensitivity), measurements were analysed at specified stress or strains. For stress-relaxation the area under the curve was calculated using numerical integration and used to compare control and PTOA ACL properties.

Normality and statistical analysis were calculated using GraphPad Prism (GraphPad v8) (GraphPad Software, CA, USA). Data normality was assessed with Shapiro–Wilk tests. For ACL CSA and length measurements, an unpaired t-test was used to compare between control and test groups. Data was not normally distributed for the stress-strain of the PTOA and control group, and therefore, a nonparametric statistical test was performed (Mann-Whitney test). A nonparametric Friedman test was used when comparing the stress-strain at the three different strain rates, followed by a Dunn’s multiple comparison test. All other ACL properties showed normal distribution and an unpaired t-test (two tailed) was performed to compare between control and test groups. A repeated measures ANOVA was used when comparing the tangent modulus-stress and hysteresis at different strain rates, followed by a Bonferroni post-hoc test. For stress-relaxation, when comparing the area under the curve of the normalised tangent modulus of the 1%/s and the 10%/s strain rates, a paired t-test (two-tailed) was used. A Friedman test, repeated measures ANOVA and paired t-test were possible since measurements were taken from the same ACL at different strain rates. Significance was set at p < 0.05.

## 4 Results

### 4.1 Knee joint space mineralisation increased with PTOA disease progression

Calcified and ossified tissues in the murine knee joint was visualized and quantified using μCT imaging. The non-invasive PTOA knee joint had increased mineralisation compared to healthy control knee joints at both 4-(33% increase, p<0.01) and 14-weeks (46% increase, p<0.001) following trauma (Fig. 2A). Mineralisation also significantly increased at 14 weeks post-trauma compared to 4 weeks post-trauma (p=0.02) (Fig. 2A), suggestive of progressive OA pathology. Furthermore, representative reconstructed 3D μCT images demonstrated the mineralisation was localised primarily to the lateral and posterior menisci and meniscal ligaments (Fig. 2B) and these increases in knee joint space mineralisation were indicate structural changes to knee joint tissues following trauma.

**Figure 2:**
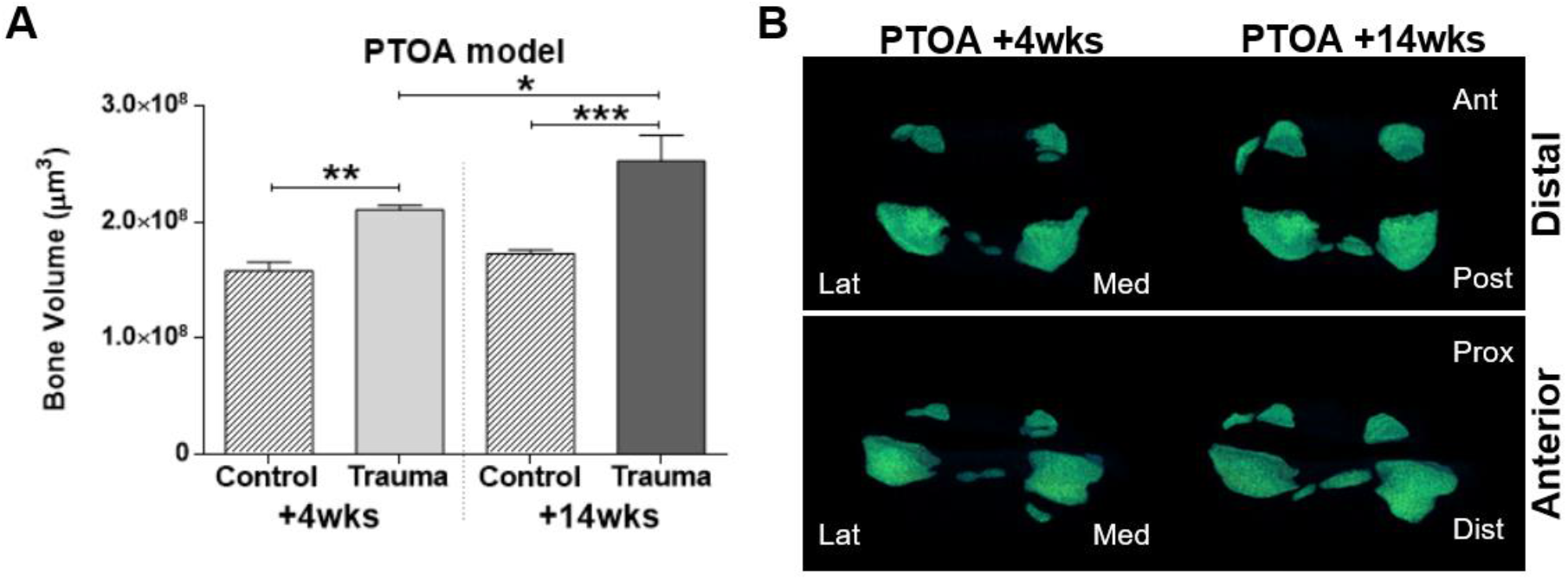
Quantification and 3D images of joint space mineralisation in post-traumatic OA (PTOA) murine knee joints using μCT analysis. A) The volume of mineralised tissue in the joint space increased between the 4 and 14 weeks timepoints of post-traumatic OA (following 2 weeks of repetitive trauma loading) compared to non-loaded healthy controls. B) 3D enlargement of the meniscus and ectopic mineralisation nodules showed mineralisation in the lateral and posterior compartments of the knee joint.

### 4.2 Histological assessment of the PTOA ACL revealed cellular hypertrophy and expression of chondrogenic markers

Histological staining of the knee joint revealed differences in structural and cellular composition of healthy and PTOA knee joints ligaments. In the healthy control knee joints, the ACL midsubstance comprised of spindle-shaped cells with aligned fibres that stain only weakly with TB (Fig. 3A) and green collagen birefringence (Fig. 3E). In the healthy ACL-tibial enthesis region, there were fibrocartilaginous cells with rounded morphology surrounded by TB staining (Fig. 3A, black arrow) and red collagen birefringence (Fig. 3E). In the PTOA knee joint, the ACL midsubstance had consistent stronger TB staining and cells with rounded cell morphology (Fig. 3B, yellow arrow). Furthermore, in the PTOA ACL-tibial enthesis there was a distinct TB region encompassing cells with rounded morphology (Fig. 3B, black arrow), which corresponded with red collagen birefringence that extended towards the midbody region (Fig. 3F, blue arrows). Increases in TB staining and red collagen birefringence in the diseased ACL indicate a change in ECM composition and organisation with disease.

**Figure 3:**
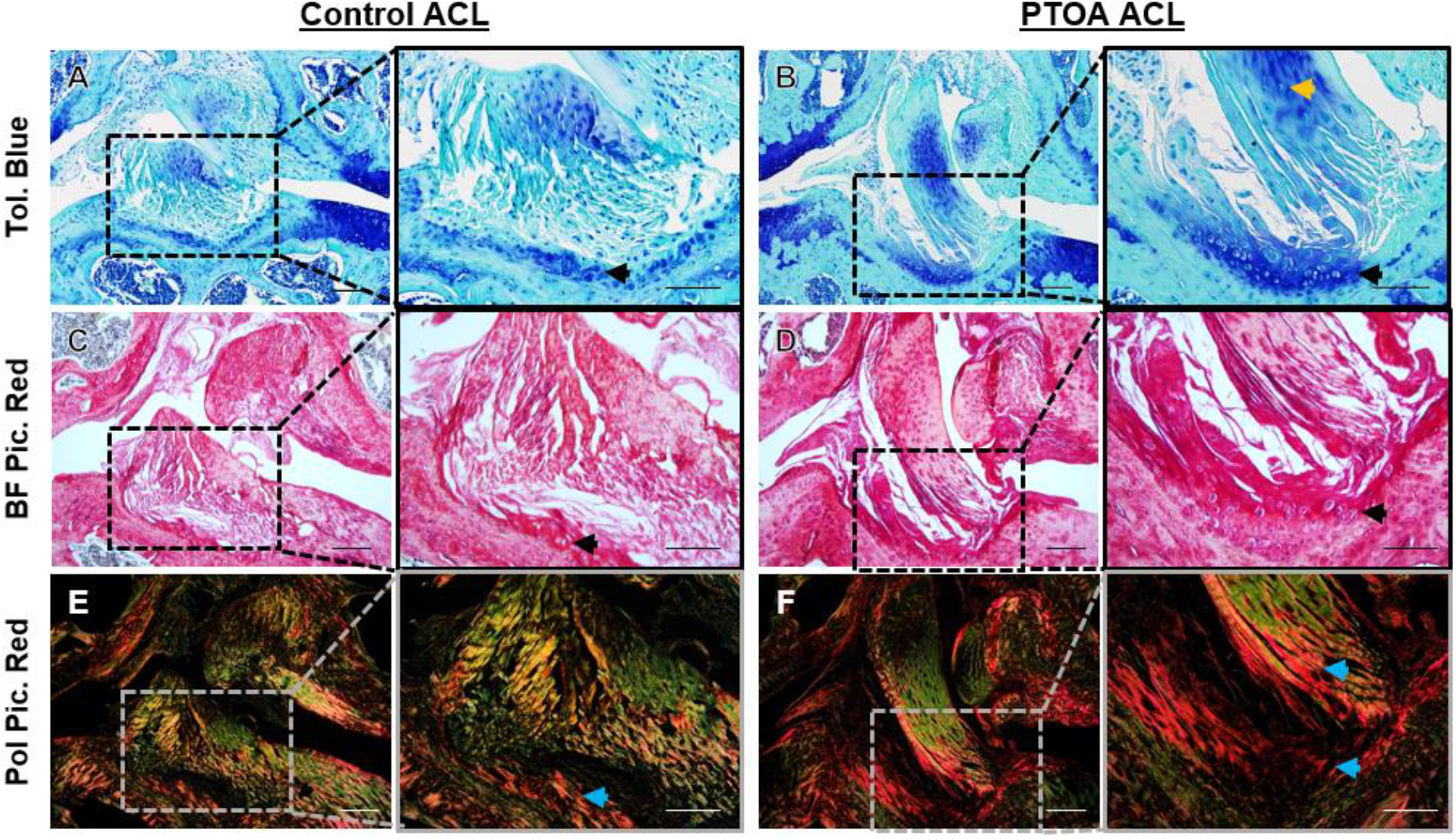
Representative histological staining in the anterior cruciate ligament (ACL) of control and post-traumatic OA (PTOA) murine knee joints. A-B) Toluidine blue (Tol. Blue) staining of healthy control ACL and PTOA ACL showed toluidine blue staining in the midsubstance and in the tibial enthesis of the PTOA ACL (black arrow). C-D) Brightfield Picrosirius Red (BF Pic. Red) staining confirmed rounded-cell morphology in the tibial enthesis of the PTOA ACL (black arrow). E-F) Polarised Picrosirius Red (Pol Pic. Red) showed red collagen birefringence in the tibial enthesis extending towards the midsubstance of the PTOA ACL (yellow arrows). Scale is 100um and 50um for the low and higher magnification images, respectively.

To further understand these ECM changes, staining for specific matrix and cellular markers, via immunohistochemistry, confirmed modifications to ECM composition with PTOA. Collagen type II (COL2) was present as expected in the fibrocartilaginous region of the ACL-tibial enthesis of the healthy control (Fig. 4C, black arrow). In the PTOA knee joint, COL2 staining expanded from the ACL-tibial enthesis (Fig. 4D) towards the ACL midsubstance region (Fig. 4E, black arrow). Cellular changes were also confirmed in the PTOA knee joint and included expression of chondrogenic marker SOX9, hypertrophic marker RUNX2, and small-leucine rich proteoglycan (SLRP) Asporin (ASPN). Both SOX9 and RUNX2 expression was faint in the healthy control ACL-tibial enthesis (Fig. 4F, I), but were clearly expressed in cells at the ACL-tibial enthesis (Fig. 4G, J) and ACL midsubstance (Fig. 4H, K) in the PTOA knee joint. Similarly, pericellular ASPN expression was found in the healthy control knee joint in the ACL-tibial enthesis in limited cells (Fig. 4L), but ASPN expression was found throughout the ACL (black arrows) and in surrounding regions (red arrow) (Fig. 4M-N) in the PTOA knee joint.

**Figure 4:**
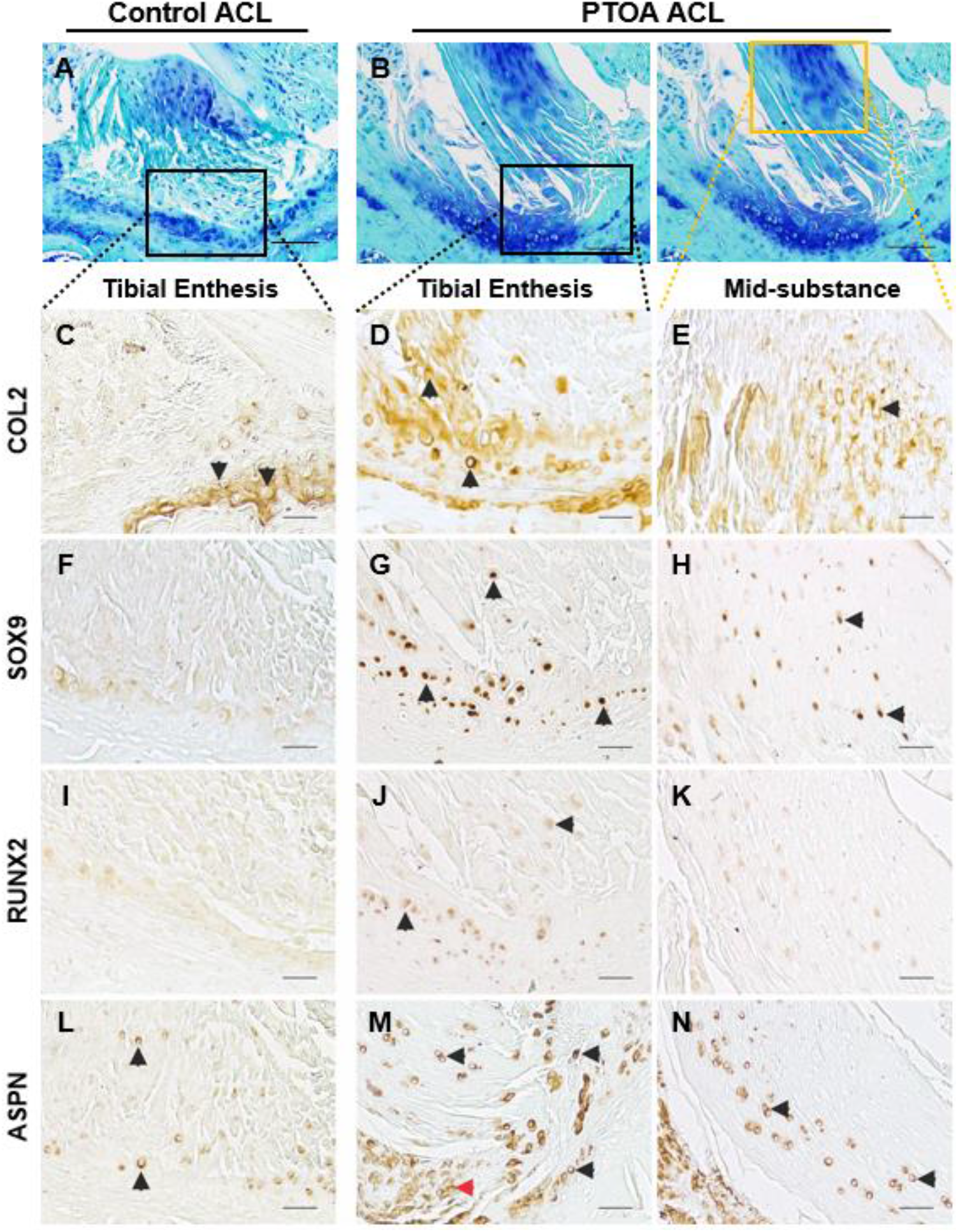
Representative marker expression in the anterior cruciate ligament (ACL) of control and post-traumatic OA (PTOA) murine knee joints with immunohistochemistry. A-B) The ACLs of healthy control knee joints and PTOA knee joints (B) where analysed at the tibial enthesis (black box) and midbody (yellow box) regions. C-E) Collagen type II (COL2) expression was present in the fibrocartilaginous tibial enthesis of the healthy control ACL and in the tibial enthesis and mid-body regions of the PTOA ACL (black arrows). F-H) SOX9 chondrogenesis transcriptional factor expression was also found in the tibial enthesis and mid-body regions of the PTOA ACL (black arrows). I-K) RUNX2 hypertrophic transcriptional factor was expressed in the tibial enthesis region of the PTOA ACL (black arrows). L-N) Asporin (ASPN), a small leucine-rich proteoglycan, was expression in the tibial enthesis of the control ACL and extended within the mid-substance of the PTOA ACL. Scale is 25um for immunohistochemistry (higher magnification), and 50um for A-B (lower magnification).

Immunohistochemistry confirmed changes to the ECM composition and cellular phenotype in diseased mouse ACL, with strong suggestions of early endochondral ossification pathways being activated (SOX9, COL2, ASPN).

### 4.3 Viscoelastic stiffness decreased, strain rate sensitivity decreased, and relaxation increased in the murine PTOA knee joint ACL

Mechanical properties of murine ACLs following PTOA development were assessed and compared to healthy controls. Stress-strain plots of behaviour at strain rates of 0.1%/s, 1%/s and 10%/s showed exponential viscoelastic behaviour in healthy control ACLs (Fig. 5A). PTOA ACLs also demonstrated similar viscoelastic behaviour and stress was compared to healthy control ACLs at several strains. At 0.03 mm/mm strain, the midpoint, stress in the PTOA ACLs was not significantly different from the control ACLs at any strain rate (p=0.19 for 0.1%/s, p=0.20 for 1%/s, p=0.21 for 10%/s) (Fig. 5A). However, the tangent modulus-stress plots, which provide stiffness estimates at different stress levels, demonstrated reductions in the tangent modulus in diseased ACLs. At 2MPa stress, the midpoint, the tangent modulus was significantly decreased by 20% at 0.1%/s strain rate relative to control ACLs (p=0.03), 20% at 1%/s strain rate (p=0.02), and 21% at 10%/s strain rate (p=0.02) (Fig. 5B). These results suggest a decrease in ACL stiffness in the toe-region of the PTOA knee joint stress-strain behaviour compared to the healthy control ACL.

**Figure 5:**
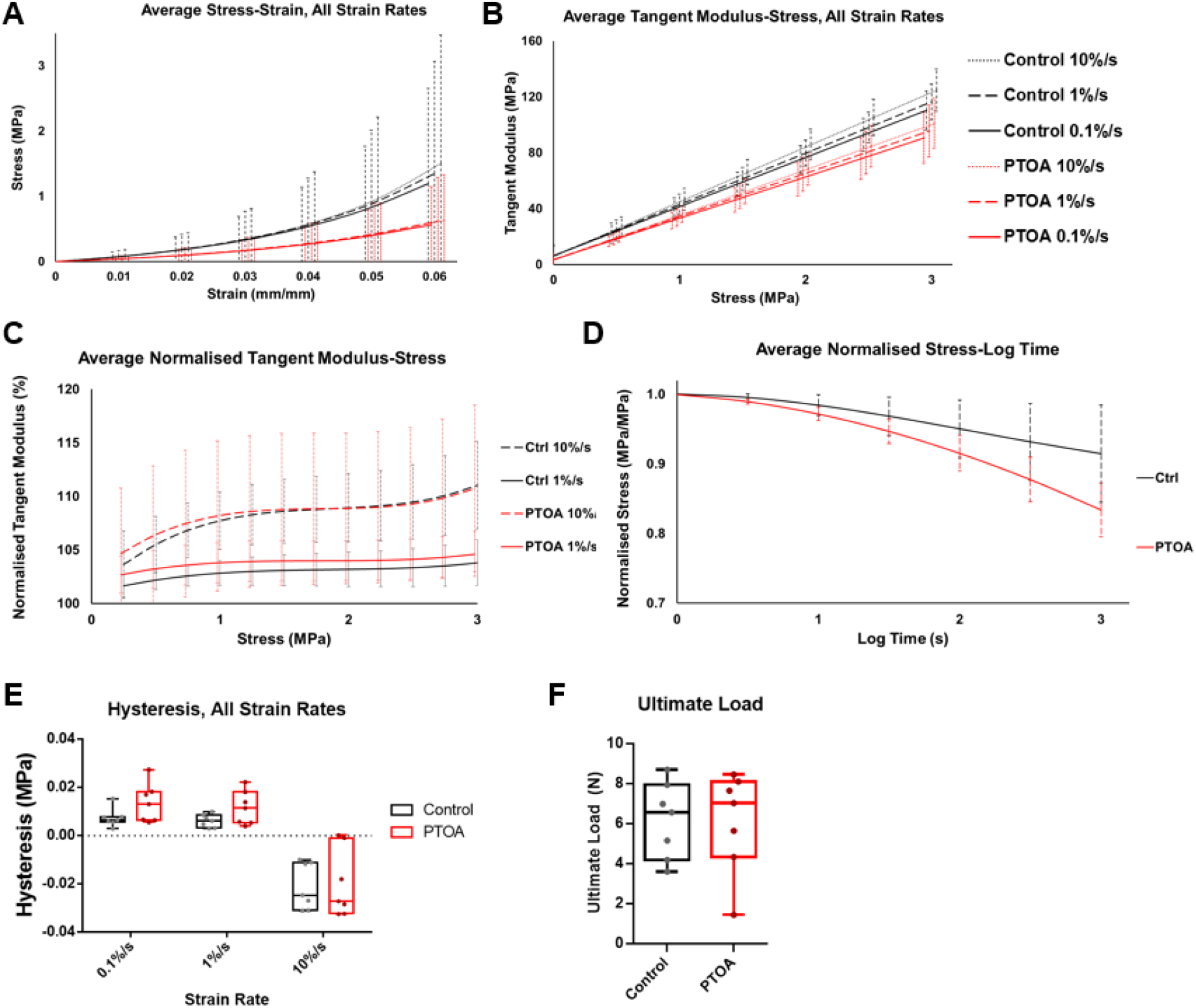
Mechanical and viscoelastic properties of the anterior cruciate ligament (ACL) of control and post-traumatic OA (PTOA) murine knee joints. A-B) Average stress-strain and tangent modulus-stress curves showed a decrease in stiffness at all strain rates for the PTOA ACLs compared to the control ACLs. C) Normalised tangent modulus-stress curves (normalised from the 0.1%/s strain rate curve) compared differences in 1%/s and 10%/s strain rates, which were statistically significant in the control ACLs (p<0.01) but not in the PTOA ACLs (0=0.07), suggesting a lack of strain rate sensitivity between 1%/s and 10%/s of the PTOA ACL. D) Stress-relaxation curves showed lower normalised stress in the PTOA ACLs. E) Hysteresis curves at different strain rates showed no statistical difference between control and PTOA (0.1%/s: p=0.09, 1%/s: p=0.08, 10%/s: p=0.88). F) Ultimate load at failure showed no statistical difference between control and PTOA ACLs. Legend: Control = black, PTOA = red.

Strain-rate sensitivity was also analysed in the stress-strain, tangent modulus-stress, and normalised tangent modulus-stress curves. In the stress-strain curves, at 0.03 mm/mm strain, no significant differences were found between different strain rates in the control ACLs (p=0.17) and the PTOA ACLs (p=0.28) (Fig. 5A). In the tangent modulus-stress curves, strain-rate differences were statistically significant at 2 MPa stress for both the control (p<0.01) and the PTOA ACLs (p<0.01) (Fig. 5B), suggestive of strain rate sensitivity in both ACL groups. However, post-hoc analysis comparing specific strain rates revealed differences at higher strain rates. In the PTOA ACLs, at 2 MPa of stress, there was no statistical differences between the 0.1%/s and 1%/s strain rates and the 1%/s and 10%/s strain rates (p=0.15 for both). In the control ACLs, at 2 MPa, post-hoc analysis showed statistical differences between all strain rates (p<0.01 for all). This would suggest a decrease in strain rate sensitivity in the PTOA ACLs. Normalised tangent modulus-stress curves confirmed these findings as strain rate differences between 1%/s and 10%/s led to significant differences in results at 2MPa of stress in the control ACLs (p<0.01) but not in the PTOA ACLs (p=0.07) (Fig. 5C) suggestive of a decrease in strain rate sensitivity in the PTOA ACLs at higher strain rates.

The normalised stress in stress-relaxation tests was lower throughout the stress-relaxation response in the diseased ACLs compared to the healthy control ACLs (p=0.04) (Fig. 5D). This was indicative of an increased relaxation response of the PTOA ACLs compared to the healthy ACLs.

When comparing the hysteresis response between the control and PTOA ACLs, there were no statistical differences at 0.1%/s (p=0.09), 1%/s (p=0.08) and 10%/s (p=0.88) (Fig. 5E). Furthermore, negative hysteresis was found in the 10%/s strain rate for both the control and PTOA ACLs (Fig. 5E). Therefore, changes in viscoelastic behaviour were seen in the stress-relaxation but not hysteresis.

The material strength of the ACL was also measured with the ultimate load to failure, which was similar in the healthy control and PTOA ACLs. The ultimate load was 6.17 ±1.91 N and 6.10 ±2.51 N for the control and PTOA ACLs respectively (p=0.96) (Fig. 5F). This would imply similar material strengths of the control and PTOA ACLs, despite differences in viscoelastic properties.

Overall, viscoelastic properties of the PTOA ACL demonstrated a decrease in stiffness, strain rate sensitivity and normalised stress during stress-relaxation, which reinforces our hypothesis that viscoelastic properties of the ACL are modified in the PTOA knee joint.

## 5 Discussion

This study found that PTOA induced non-invasively in the murine knee joint resulted in changes in the ECM components and viscoelastic mechanical properties of the murine ACL. These findings confirm that ACL pathologies do result in functional anomalies, which could in turn contribute to the progression of OA. Specifically, we have found that structural changes during PTOA, particularly proteoglycan and COL2 deposition, were associated with a decrease in ligament stiffness, a decrease in strain rate sensitivity and an increased relaxation behaviour during stress-relaxation. In particular, changes in the strain rate sensitivity of the ACL in the PTOA models is, to our knowledge, a novel finding and suggests a decrease in stiffness at higher strain rates.

Increases in joint space mineralisation measured using non-invasive imaging tools (μCT) was used in this study as an indicator of OA progression, specifically in the lateral and posterior compartments, which correlated with the location of OA development in this PTOA model (Poulet *et al.*, 2011). An increase in mineralisation in murine OA joints has been previously shown in other OA models using the same μCT-based methodologies (Ramos-Mucci *et al.*, 2020). Indeed, joint space mineralisation in the surgical model of PTOA (destabilisation of the medial meniscus, DMM) and in the spontaneous model Str/ort mouse was associated with ossification of the menisci and medial collateral ligaments, primarily in the medial compartments of the joint, corresponding with initiation of OA development in these models. These studies suggest a strong spatial relationship between articular cartilage degeneration and pathological joint tissue mineralisation and support the use of joint μCT imaging as non-invasive measures of disease progression *in vivo*. Further studies will determine the molecular and cellular basis of this pathological joint mineralisation, including in ligaments, and whether these could be targeted to slow disease development.

Changes in the ACL matrix composition during PTOA included increased TB staining, indicative of proteoglycan deposition (Sun *et al.*, 2012), red collagen birefringence and COL2 deposition. Red collagen birefringence suggests thicker collagen organisation at the ACL-tibial enthesis (Junqueira *et al.*, 1979; Lattouf *et al.*, 2014). Interestingly, these changes in collagen organisation did not result in stiffer mechanical properties of the ACL. Viscoelastic behaviour of the OA murine ACLs showed changes in the tangent modulus, strain rate sensitivity and stress-relaxation properties. These changes included a significant decrease in the tangent modulus at all strain rates, implying a decrease in viscoelastic stiffness. It is likely that ECM components are driving this viscoelastic change, including collagen type II deposition. Indeed, collagen type II molecules are generally flexible and have a lower elastic modulus (stiffness) than collagen type I molecules (Luo *et al.*, 2004) and therefore could be driving the decrease in tangent modulus. However, to truly answer this question, the role of COL2 in viscoelastic behaviour needs to be further assessed, including whether COL2 is stretched in the toe region and how it interacts with the surrounding matrix.

Stress-relaxation curves were significantly decreased in our murine osteoarthritic ACLs, indicating an increase in relaxation. Changes in viscoelastic stress-relaxation has previously been associated with glycosaminoglycan and proteoglycan depletion in tendons (Elliott *et al.*, 2003; Legerlotz *et al.*, 2013). Proteoglycan depletion is not likely in the PTOA knee joint ligaments which had an increase in TB staining, and therefore could suggest a different matrix component is driving this change in stress-relaxation.

Increases in relaxation the ACL stress-relaxation behaviour in human OA and rheumatoid knees have been reported previously (Hagena *et al.*, 1989), reinforcing similar viscoelastic mechanical pathology between murine and human OA. However, the ACL of OA patients also had a decrease in ultimate load (Hagena *et al.*, 1989), not seen in the murine PTOA ACLs. These differences could arise from differences in disease severity between our murine OA model (samples taken during active early progression of OA) whereas human OA tissues are generally already severe when collected.

Changes in strain rate sensitivity in ACLs from OA knee joints have not been previously reported. Therefore, our study shows for the first time a decrease in the strain rate sensitivity of the higher strain rates (1% and 10%/s) which suggests a decrease in the ECM stiffness at higher strain rates. Strain rate sensitivity is another viscoelastic property associated with GAG and proteoglycan composition. A reduction in strain rate sensitivity has been reported in decorin and biglycan depleted tendons (Elliott *et al.*, 2003; Robinson *et al.*, 2017; Robinson *et al.*, 2004). Computational modelling has also supported the interfibrillar viscoelastic role of proteoglycans (Redaelli *et al.*, 2003). However, the role of proteoglycans and noncollagenous components on viscoelastic behaviour is still debated. Collagen fibrils have greater elongation at higher strain rates (Clemmer *et al.*, 2010), and theoretically collagen could play a more important role at higher strain rates. Our study also reported asporin expression in diseased ACLs, however ASPN lacks GAG chains (Maccarana *et al.*, 2017) usually associated with increased viscosity (Kershaw-Young *et al.*, 2013) and therefore is unlikely to play a role in viscoelastic behaviour. Further research is needed to fully map proteoglycan and noncollagenous composition in the ACL during OA and their role in biomechanics.

Previous studies have shown that tendon fibroblasts respond to mechanical stimuli by adjusting ECM stiffness (Lavagnino *et al.*, 2003; Screen *et al.*, 2005), therefore it is possible that ligament fibroblasts are driving this ECM and viscoelastic changes. However, the mechanotransduction pathways involved are unknown. This study showed SOX9, RUNX2 and collagen type II expression in the ACL-tibial enthesis and midbody of the murine PTOA knee joint, in the same area where we identified cells with rounded morphology. These are known markers of chondrogenesis and hypertrophy, crucial for endochondral ossification. Asporin is a known tendon and ligament SLRP (Kharaz *et al.*, 2016), also found in the cartilage of human OA patients. It plays a role inhibiting TGF-β-induced expression of the *Col2a1* gene (Kizawa *et al.*, 2005) and is a negative regulator of periodontal ligament mineralisation by interacting directly with BMP-2 (Yamada *et al.*, 2007).The presence of asporin in the diseased ACL could be an attempt of ligament cells to control pathological chondrogenesis and mineralisation but the actual effects of increased asporin in ligaments remains to be explored.

It is still unclear whether ligament viscoelastic changes contribute to OA progression. Changes to the ACL mechanical function will affect joint stability, which could in turn accelerate OA pathology. It is clear though that ACL mechanics and OA are closely related. As mentioned previously, changes in ACL viscoelasticity and mechanics have also been reported in spontaneous OA models (Anderson-MacKenzie *et al.*, 1999; Quasnichka *et al.*, 2005). Furthermore, spontaneous OA models also demonstrated similar COL2 and SOX9 expression (Ramos-Mucci *et al.*, 2020), suggesting similar pathologies in both types of OA. COL2 in ligaments and tendons has been shown to be a result of adaptation to compressive load (Benjamin and Ralphs, 1998; Buckley *et al.*, 2013), and therefore could be a direct result of the non-invasive loading regime. In addition, COL2 has also been found in human OA ligaments (Hasegawa *et al.*, 2013) and guinea pig OA ligaments (Quasnichka *et al.*, 2006). Interestingly, it has been shown that changes in the matrix stress-relaxation properties can regulate scaffold remodelling and bone formation (Darnell et al., 2017). Therefore, similar viscoelastic changes in the ligaments could be driving the chondrogenesis seen in OA ligaments. In this case, changes to the ECM mechanical properties could be driving more pathology directly.

Limitations of this study arise from the mechanical testing of the murine ACL. Stress and strain were calculated with the assumption of similar ACL length and CSA within groups, which could affect the accuracy of the stress-strain curve. Most other studies on murine ACL mechanics measured the CSA and length for each individual sample (Anderson-MacKenzie *et al.*, 1999; Connizzo *et al.*, 2015; Sun *et al.*, 2015). The small size of the murine ACL and these inaccuracies could account for the high deviation seen in the stress-strain curves and the negative hysteresis measured. Negative hysteresis has been previously reported in human tendons (Peltonen *et al.*, 2013; Zelik and Franz, 2017) and attributed to testing inaccuracies. However, negative hysteresis was only measured in the faster 10%/s strain rates at areas of high stress, similar to what was reported previously in the human Achilles tendon (Peltonen *et al.*, 2013). Past studies in human skin have attributed negative hysteresis to varying viscoelastic properties of different skin layers (Lamers *et al.*, 2013). The same could potentially apply to the ACL, negative hysteresis could be a result of differences in its ligament bundles or fascicles.

Overall, this research showed that ligament matrix composition and viscoelastic behaviour change in PTOA knee joints, potentially driven by cellular expression of chondrogenic and hypertrophic markers which could be driving OA progression. Furthermore, we have confirmed our hypothesis that structural changes in the ACL resulted in a reduction in viscoelastic stiffness and other viscoelastic properties of the ACL. This confirms that the ligament biomechanical microenvironment is altered during OA. Understanding the relationship between the structure and mechanical function is necessary to better understand the underlying OA pathways. In the future, targeting these molecular and cellular changes might improve ligament function and prevent further OA progression.

## Acknowledgments

We thank John Marrin for his assistance in collecting and digitising the biplanar X-ray data.

## Author contribution

Study concept and design: B.P., E.C., A.Elsheikh.

Acquisition, analysis and interpretation of data: all authors.

Statistical analysis: B.P., L.R., A. Eliasy.

Drafting of the manuscript: all authors.

Critical revision of the manuscript: all authors.

## 7 Supplementary material

### 7.1 Gait analysis of murine knee flexion

Gait analysis tests to determine the physiological range of motion in the murine knee joint was conducted on healthy B6CBAF1 mice (n=4) (Charles River). Biplanar X-ray was recorded using two independent 60 kW Epsilon X-ray generators (EMD Technologies, CA), 16 inch image intensifier tubes (Thales, FR), X-Ray Tubes (Varian, USA) and Phantom Miro M120 video cameras (Vision Research, USA). X-ray images of gait (Sup. Fig. 1A) were analysed using XMALab Software (Brown University, USA) and MatLab (MathWorks, R2018a Version 9.4, USA). For each mouse, two gait trials were analysed. X-ray generator pulsed radiographic exposures at 120 frames per second, and video was recorded for approximately 3 seconds. Video files were corrected for distortion, calibrated and digitised in XMALab software. Mouse gait was tracked at every frame at the following points: left hip, left knee, left ankle, right hip, right knee, right ankle, and occiput (Sup. Fig. 1B). The resulting locations of each point were exported and in MatLab a 3D model was created (Sup. Fig. 1C) and knee speed and knee joint flexion was measured at every frame. The results from this gait analysis of physiological murine knee flexion were used to determine the ideal physiological angle for mechanical testing of the femur-ACL-tibia complex.

**Supplemental Figure 1:**
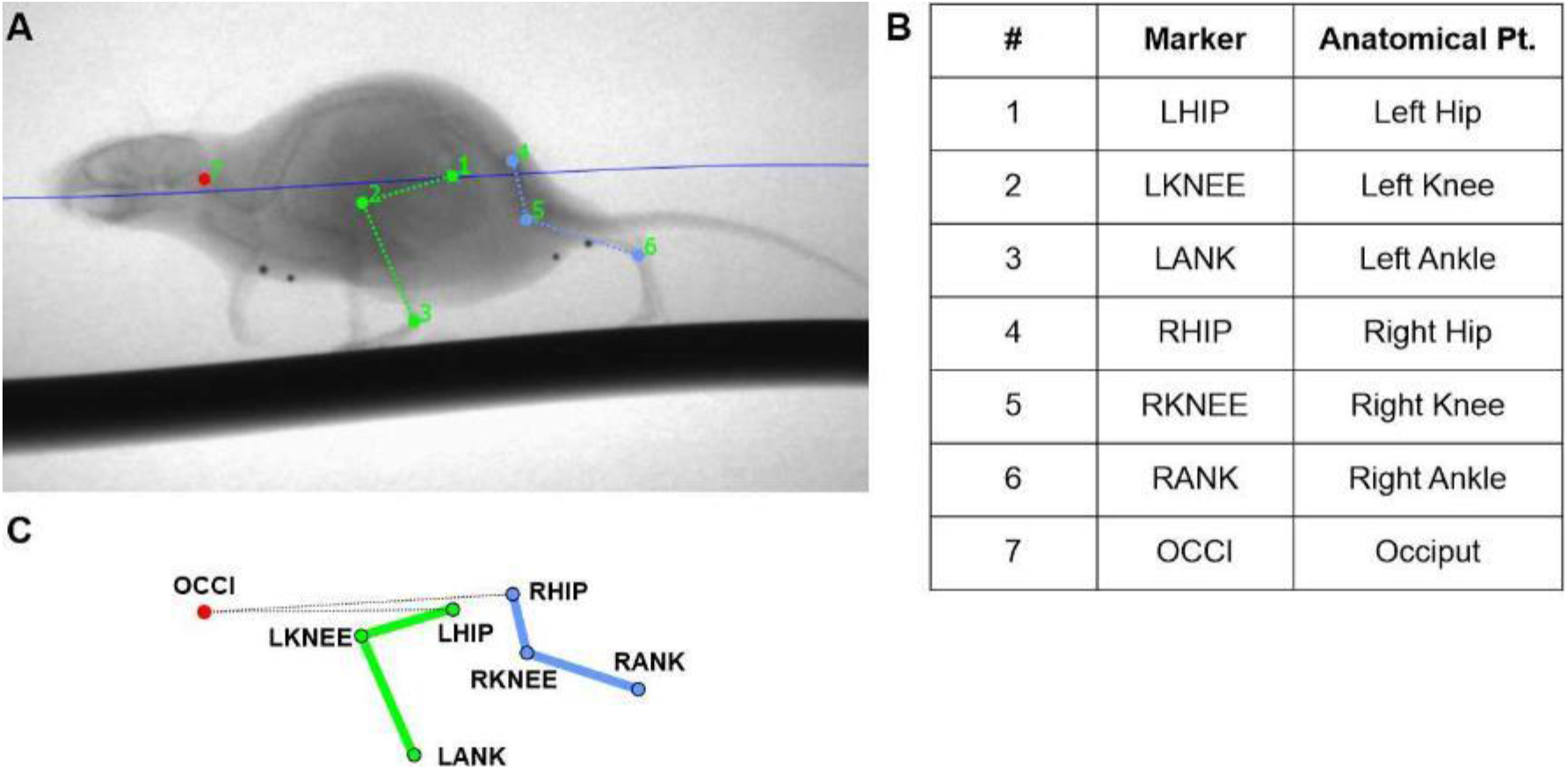
Murine knee flexion measurements using biplanar X-ray. A) X-ray images were imported to XMALab software (Brown University, USA) where each anatomical point (Pt.) was tracked. B) Anatomical points tracked were the following: left hip (1, LHIP), left knee (2, LKNEE), left ankle (3, LANK), right hip (4, RHIP), right knee (5, RKNEE), right ankle (6, RANK), and occiput (7, OCCI). C) A 3D model of murine gait for the left and right hind-legs was created from which physiological knee range of motion during murine gait was calculated.

Maximum knee flexion was 100.8° in the left knee joints and 100.5° in the right knee joint (Sup. Table 1). Minimum knee flexion was 56.5° and 55.6° in the left and right knee respectively (Sup. Table 1). Maximum and minimum angles of knee flexion in both the left and right knee joints of the B6CBAF1 (n=4) mice were similar. This data suggests that the physiological range of murine knee flexion during gait was between 55-100°. In order to remain within the physiological range, mechanical viscoelastic and material testing of the murine ACL was performed at 90° of knee flexion, because it allowed for uniform tensile testing along the axis of the ligament, similar to a previous study by Warden et al (Warden *et al.*, 2006).

**Supplemental Table 1.** Angles of murine knee joint flexion and gait speed as determined by biplanar radiography. The average minimum (Min) and maximum (Max) knee flexion was calculated in degrees for both the left and right murine knee joint. Knee flexion ranged from 56.5 to 100.8° in the left knee, and 55.6 and 100.5° in the right knee joint. Relative standard deviation (RSD) ranged from 6.7% to 13.3%.

|  | Degrees (°) |  |  |  | Gait speed (mm/s) |  |
| --- | --- | --- | --- | --- | --- | --- |
|  | Min<br>Left Knee | Max<br>Left Knee | Min<br>Right Knee | Max<br>Right Knee | Left knee | Right knee |
| **Avg.** | 56.5 | 100.8 | 55.6 | 100.5 | 135.6 | 138.7 |
| **RSD (%)** | 8.8 | 13.3 | 6.7 | 11.5 | 24.3 | 19.1 |

### 7.2 Anterior cruciate ligament (ACL) measurements of PTOA knee joints

ACL length and CSA were measured using μCT imaging. The ACL length of the control and PTOA knee joint ACLs was 1.14 ±0.10 mm and 1.09 ±0.09 mm respectively and were not significantly different (p=0.08). The ACL CSA was 0.105 ±0.017 mm^2^ for the control knee joint and 0.109 ±0.017 mm^2^ for the PTOA knee joint and were not significantly different (p=0.6) (Sup. Table 2). The ACL measurements for each murine group were used to calculate their viscoelastic properties.

**Supplemental Table 2.** Murine knee joint anterior cruciate ligament (ACL) measurements. Measurements of the ACL were taken from healthy and PTOA knee joints and included ACL length and ACL cross-sectional area (CSA). For each measurement relative standard deviation (RSD) and percent different (%diff) was calculated. CSA and length of control and PTOA ACLs were not significantly different (p=0.6 for CSA, and p=0.08 for length).

| Group | ACL Length |  |  | ACL CSA |  |  |
| --- | --- | --- | --- | --- | --- | --- |
|  | Length (mm) | RSD (%) | %diff | CSA (mm<sup>2</sup>) | RSD (%) | %diff |
| Control Knee | 1.1441 | 9.1 | 5.12 | 0.1046 | 16.2 | 3.66 |
| PTOA Knee | 1.0869 | 8.4 |  | 0.1085 | 16.1 |  |

